# Homology-Based Enzymatic Assembly of Modular T7 Phage Genome

**DOI:** 10.1101/2025.04.24.650493

**Authors:** Julia Vu, Zane Chan, Nicholas Murphy, Akio Shirali, Katie Han, Nils JH Averesch, Phillip Kyriakakis

## Abstract

Bacteriophages, viruses that infect bacteria, are pivotal in therapeutic, industrial, and bio-detection applications due to their unique ability to inject DNA into bacterial hosts and change the genetics and behavior of whole bacterial populations. Genetic engineering of phage genomes has expanded their potential applications with several established methods such as Golden Gate assembly, yeast cloning, λ Red recombineering, and CRISPR-Cas systems. Here, we present a novel, efficient method to design and make synthetic bacteriophages *in vitro* without using restriction enzymes to allow for modular insertion of DNA fragments into the phage genome. The ability to create synthetic bacteriophages *in vitro* without the use of restriction enzymes allows for simpler engineering without the hassle or cloning limitations encountered when building domesticated bacteriophage genomes.

## Introduction

Bacteriophages, viruses that infect and replicate in bacteria, are the most abundant and diverse biological entities on Earth, with an estimated >10^30^ particles in our biosphere (Chibani-Chennoufi *et al*. 2004; Czajkowski 2019; Mann 2005). The bacteriophage’s ability to alter bacterial behavior at the population scale makes them valuable for therapeutic (Chan & Abedon 2015; Dedrick *et al*. 2019; Kortright *et al*. 2019), industrial (Ji *et al*. 2021; Kawacka *et* al. 2020), and bio-detection applications (Al-Hindi *et al*. 2022; Kim *et al*. 2021; Pearson *et al*. 1996). Genetic engineering techniques have enabled facile modification of phage genomes, expanding their native functions and enhancing their potential applications (Jia et al. 2023).

To enable the production and optimization of these tools, several approaches to phage genetic engineering have been developed. *In vivo* cloning methods such as λ Red recombineering (Jensen *et al*. 2020; Oppenheim *et al*. 2004; Thomason *et al*. 2023) and CRISPR-Cas systems (Kiro *et al*. 2014; Lemay *et al*. 2017; Martel & Moineau 2014) facilitate the editing of the intact phage genome. More recently, phage genomes were inserted into a Yeast Artificial Chromosome (YAC) and edited in yeast (Ando *et al*. 2015). This method allows for genome assembly by transforming fragments containing sequence overlaps into the YAC along with a selection marker. Once encoded in the YAC, the phage genome can be edited through homologous recombination using a removable selection marker such as URA3. In bacteria, *in vivo* approaches are limited by the need for a selection method and CRISPR-immunity mechanisms in phages (Bondy-Denomy *et al*. 2013; Hossain *et al*. 2021). Additionally, the process of combining, modifying, and then isolating phage genomes before finally rebooting the genome is time-consuming and laborious.

In contrast, *in vitro* phage genome assembly strategies permit a more rapid assembly and rebooting pipeline. There are several *in vitro* phage genome assembly methods, including polymerase cycling (Smith *et al*. 2003), Golden Gate Assembly (Potapov *et al*. 2018; Pryor *et al*. 2022), isothermal recombination (iPac) (Nozaki 2022), and PHage Engineering by *in vitro* Gene Expression and Selection (PHEIGES) (Levrier *et al*. 2024). Each has different limitations, but they allow direct rebooting of the synthetic phages after genome assembly. For example, Golden Gate requires domestication (removal of specific restriction sites), iPac requires a packaging kit (Nozaki 2022), and PHEIGES requires a TXTL kit (Levrier *et al*. 2024). Using Gibson Assembly/NEBuilder to construct the genomes in vitro is cheaper and easier than iPAC and TXTL and does not require domestication like Golden Gate (Pulkkinen *et al*. 2019). The capability to assemble without domestication is crucial for engineering bacteriophage because promoters cannot be modified by using alternative codons, as with protein-coding genes, and the structures of transcribed RNAs can have functions that would be interfered with by codon optimization (Mortimer *et al*. 2014).

One of these limitations arises when attempting to test multiple versions of a small segment of the genome. While previous approaches allow for the reassembly of many genome fragments along with a single varied fragment, this process requires reassembling five or more fragments with high fidelity each time. Additionally, testing many versions using a TXTL or packaging kit can quickly become prohibitively expensive. Another approach would be to modularize the T7 bacteriophage genome for simple insertions of DNA parts into a specific location in the genome using an engineered “landing pad” for a single new fragment. However, producing a phage with a landing pad in some genomic regions can affect the fitness of the phage or make it completely inviable. For example, T7 bacteriophage control systems often reside in promoter elements (*e*.*g*., at a ribosome binding site) that cannot be modified without altering function (Weisberg et al. 2008). Alternatively, designing synthetic bacteriophages without using restriction enzymes allows for engineering without the hassle or cloning limitations encountered when building “domesticated” bacteriophage genomes (Ellis et al. 2011; Kilcher & Loessner 2019). Leveraging *in vitro* homology-based assembly methods, such as Gibson Assembly, for phage genome reconstruction allows for seamless assembly that does not need a Type IIS restriction enzyme to generate the “overhangs” required with Golden Gate Assembly (Gibson et al. 2009; Yeom et al. 2020). This approach allows for optimizing the homology regions used for assembly or for optimizing the position of the landing pad. In addition, using Gibson Assembly enables the insertion of Type IIS restriction sites that would otherwise be cut during Golden Gate assembly.

In this work, we employed a hybrid approach where we domesticated the genome to remove SapI sites so they could be used in our landing pad, but assembled the genome using Gibson rather than Golden Gate. This allowed us to reintroduce the SapI site into the genome during assembly. In addition to allowing the use of SapI in our landing pad, engineering it without SapI also made it easy to distinguish from the wild-type (WT) phage. In our first design-test-build-learn cycle, we found that inserting some sequences into the promoter region of gene 2.5 indeed affected the fitness of the phage. Analysis of the structure of the WT and modified gene 2.5 RNA transcripts provided insights into the factors contributing to the low viability and informed the design of a new landing pad with minimal effects on the fitness. Since gene 2.5 is essential, this approach could facilitate phage library testing to screen for riboregulators that respond to RNA triggers (Goicoechea Serrano et al. 2024) and for performing directed evolution to make these regulators more robust. Moreover, this strategy could be applied more broadly to insert DNA libraries into arbitrary sites with high efficiency.

## Results and Discussion

### Modified Phage Genome Design

We sought to develop a system for T7 genome assembly that is modular and serves as a streamlined platform for inserting and testing gene fragments into a specific region of the genome **(Figure 1A)**.

**Figure 1:**
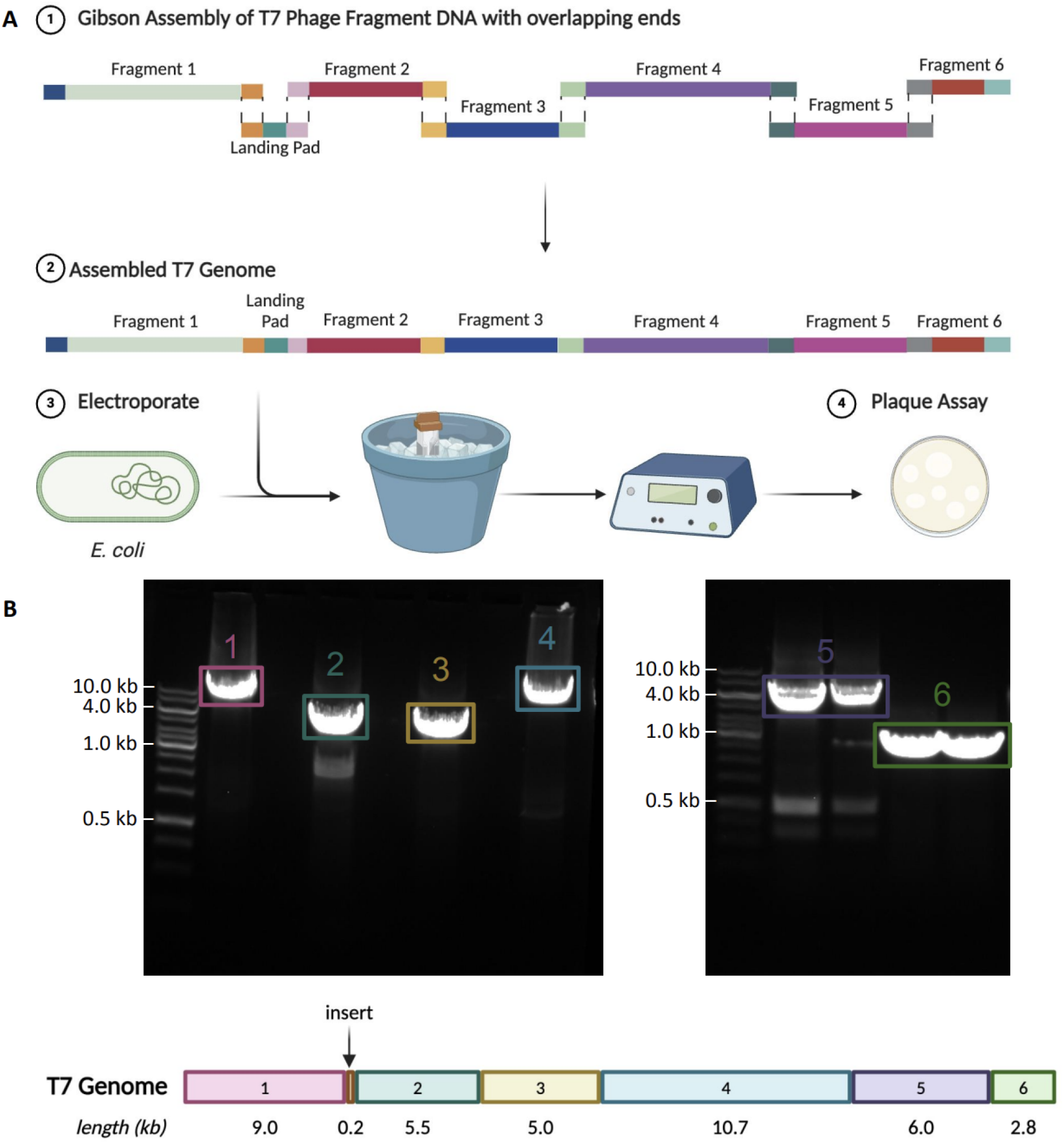
Homology-based enzymatic assembly of the T7 phage genome. **(A)** An overview of the genome assembly procedure with the genome fragments and overlaps annotated. The “landing pad” is an engineered cloning site for inserting regulatory elements for gene 2.5. **(B)** PCR results showing the amplification of each genome fragment.

To test our assembly method, we engineered a modified T7 bacteriophage containing a “genetic landing pad” upstream of the essential T7 gene, “gene 2.5”. This gene encodes a single-stranded DNA-binding protein that interacts with T7 gene 5 (DNA polymerase) and gene 4 (helicase and primase) proteins, which are crucial for T7 DNA synthesis (Kim & Richardson 1993). Therefore, we hypothesized that regulating this pathway by inserting a riboregulator (Isaacs et al. 2004), toehold switch (Green et al. 2014; Pardee et al. 2014), or other regulatory elements would serve as control switches of phage DNA replication. These features can make for a useful insertion site for a genetic landing pad containing unique restriction sites that allow future insertions of DNA fragment libraries into digested phage DNA. We initially designed the genome to have all SapI sites removed so that SapI could be used to cut out the landing pad for inserting control switches. This proved problematic (details below), so we redesigned the landing pad to use SapI and EcoRI sites. Fragment sizes were chosen based on the location of the native SapI sites so that we could remove them by PCR and assemble the resulting fragments using Gibson Assembly directly into a functional genome.

### Utilizing Gibson Assembly for Assembling Phage Genomes in vitro

As proof of principle for the seamless assembly of a custom phage genome without restriction enzymes, we first assembled a T7 bacteriophage free from SapI sites using Gibson Assembly. Pieces of the phage genome were amplified by PCR from a commercial WT T7 genome (Boca Scientific Inc, NC1143564), fusing homology arms to adjacent fragments. The resulting fragments are shown in **Figure 1B**.

To assess the assembly between each junction quickly, we devised a method to check for assembly *in vitro*. For rapid PCR screening and efficient amplification, we designed primers that generate small amplicons that span the junctions of the fragments **(Figure 2B)**. Once junctions were confirmed, the genome was rebooted in *E. coli*, yielding plaques, and indicating viable phage **(Figure 2A)**.

**Figure 2:**
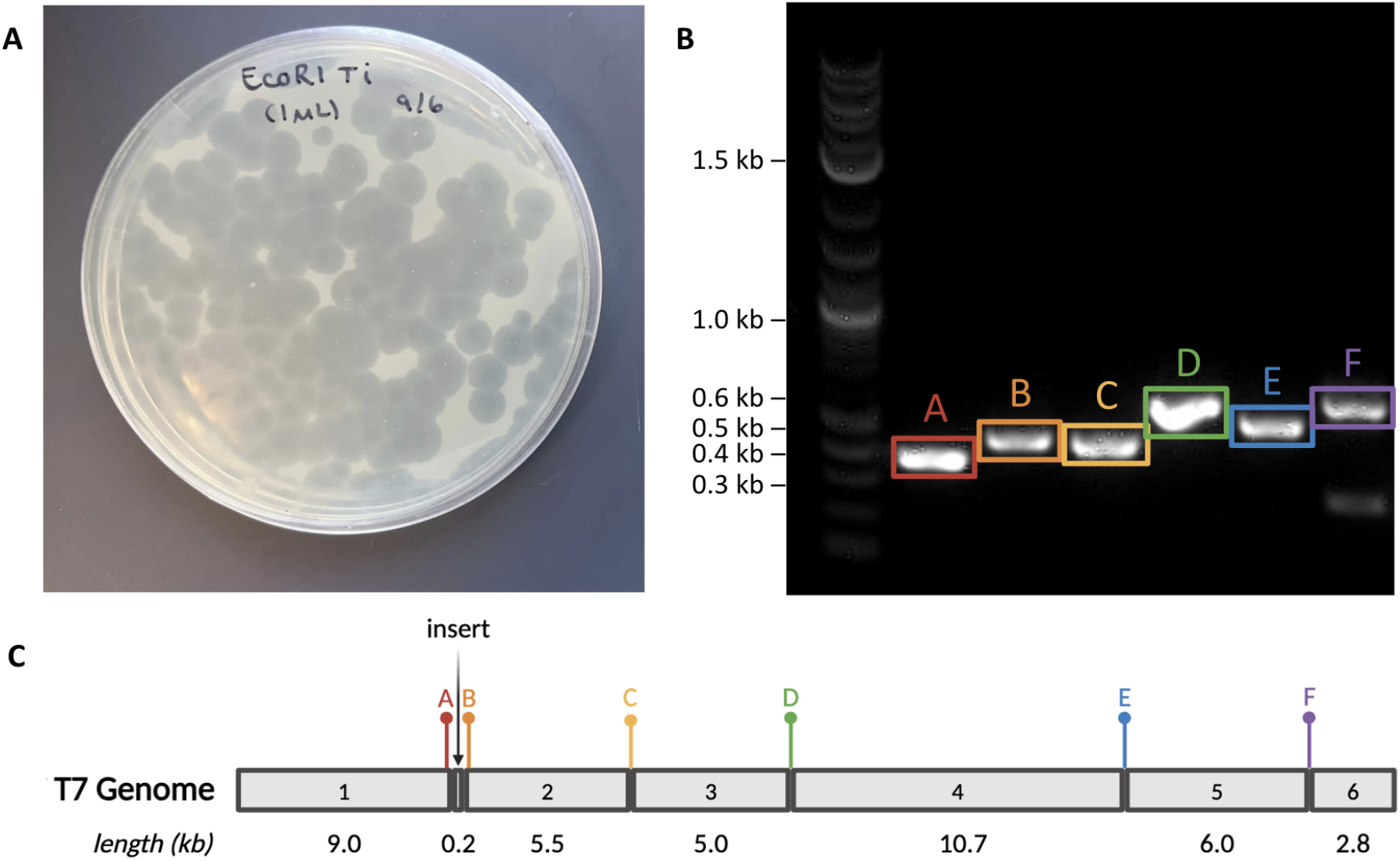
Assembled phage containing landing pad insert rebooted in DH10B. **(A)** The linear genome phage containing the Sapl and EcoRl containing landing pad transformed in DH10B. **(B)** PCR assessment of in vitro assembly. Gibson Assembly reactions were amplified via PCR using primers spanning the assembly junctions. **(C)** Genome map of linear assembled phage with location of respective junctions.

This experiment suggested that all the fragments could be assembled *in vitro*. However, it has been shown that the DNA could be assembled after transformation, so the PCR screen does not necessarily prove that the entire genome was assembled as a single fragment. To assess the assembly more thoroughly, the purified assembly was sequenced using long-read sequencing (Plasmidsaurus). As expected, the majority of DNA fragments sequenced were only partially assembled **(Figure 3)**. None of the raw reads were as long as the WT phage genome (40 kb); the longest fragment sequenced was 30.8 kb. It is also possible that some full-length genome is assembled to form our observed plaques, but were not found in our sequencing results. One explanation for why full-length genomes were not observed in the sequencing is that the DNA is partially fragmented before long-read sequencing with Plasmidsaurus. Therefore, it remains possible that the T7 genome is only partially assembled *in vitro* prior to rebooting in the bacterial host or that the mixture that is transformed contains fragments that are annealed but not covalently linked, as is the case in iPac (Nozaki 2022) and PHEIGES (Levrier et al. 2024) methods.

**Figure 3:**
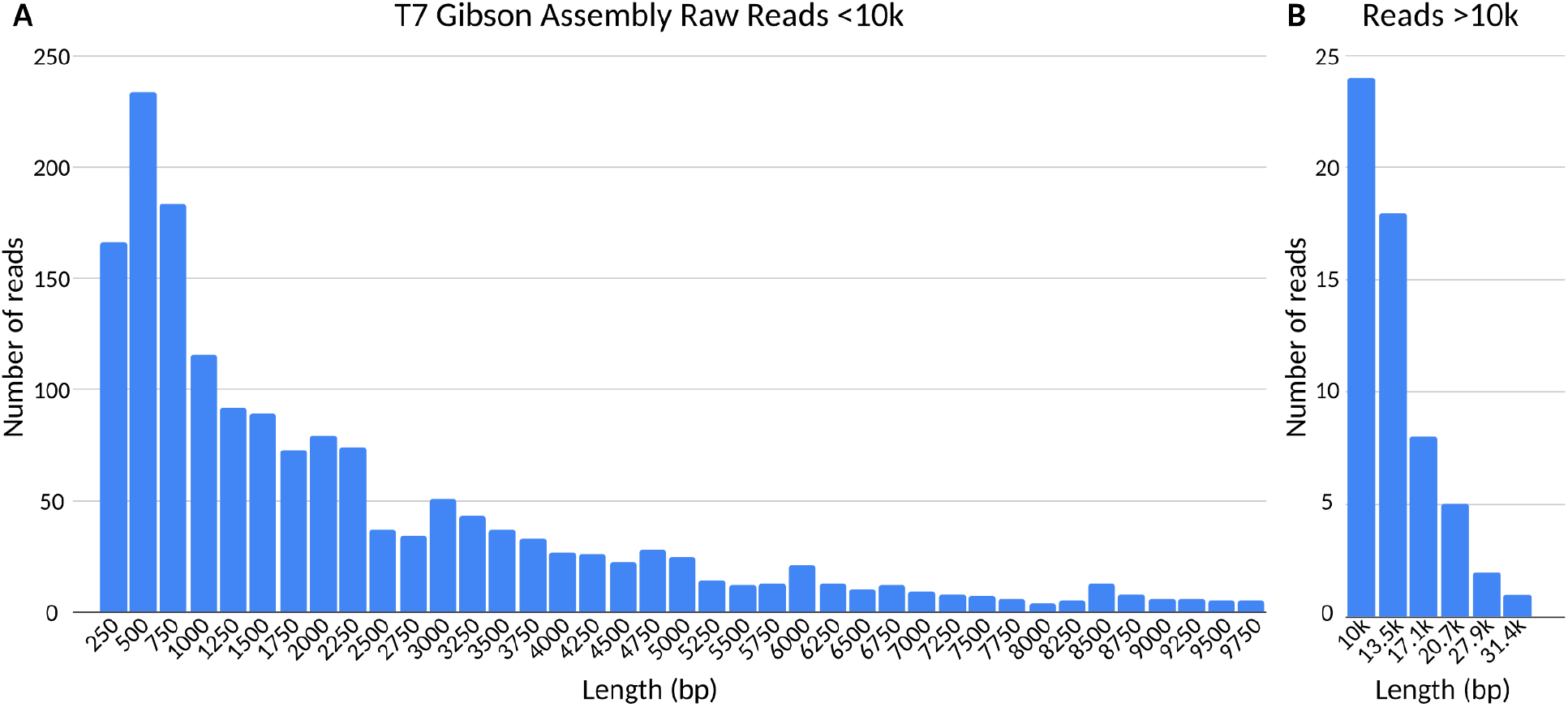
Histograms of the purified T7 Gibson Assembly reaction mixture long-read sequencing raw reads. **(A)** Raw read sizes and coverage of fragments less than lOkbp. **(B)** Raw read sizes and coverage of fragments over lOkbp.

To investigate the contribution of Gibson assembly of either partial or full-length phage genome construction, we additionally compared transforming the unassembled DNA fragments with the Gibson assembled genome. Plaque assays of T7 rebooted from Gibson-assembled and unassembled DNA fragments showed an obviously higher number of plaques when the genome was assembled **(Figure 4)**. This suggests that pre-assembly of the phage facilitates more effective rebooting, likely by full-length genome transformation or by a partial assembly, leading to more efficient *in vivo* assembly. Transforming the fragments alone had very low efficiency, which may not be very reliable, especially when assembling genomes with fitness costs. Given the efficiency observed with assembling and rebooting such a large genome from so many fragments, there is no need to package the assembled DNA into phages *in vitro* prior to introducing it into a bacterial host.

**Figure 4:**
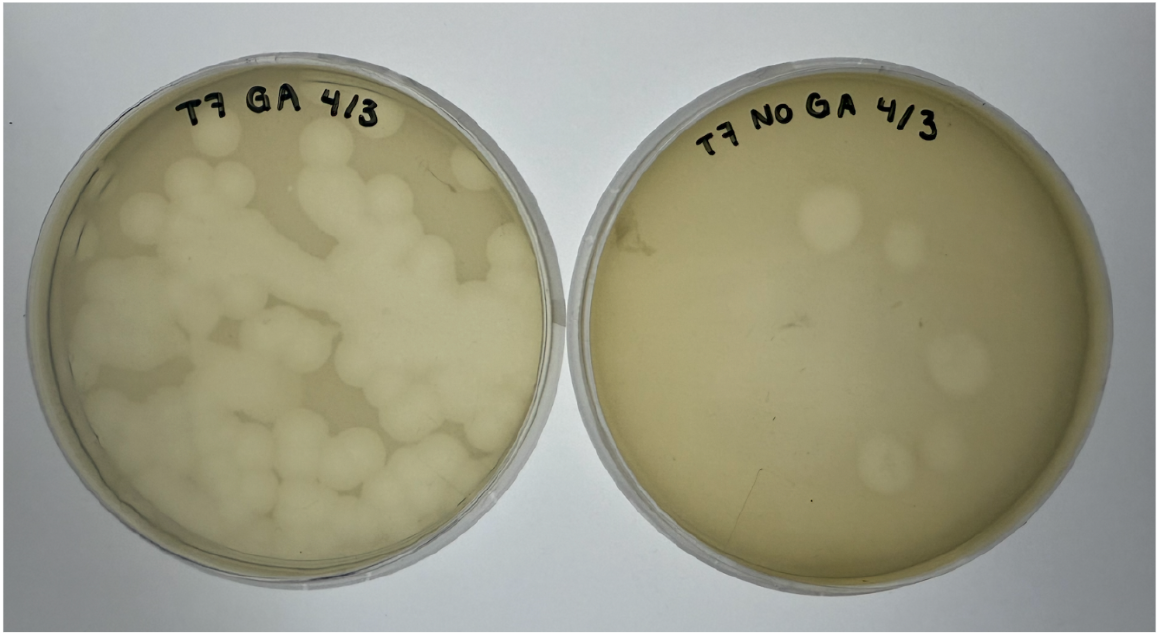
Homology-based enzymatic assembly ofT7 phage genome prior to rebooting improves plaque formation efficiency. Plaque assay of T7 phage rebooted from Gibson-assembled fragments (left) versus unassembled fragments (right)

### Circular Phage Genome Assembly

Because Gibson Assembly reaction mixtures contain an exonuclease that would digest the ends of the linear phage genome, we reasoned that assembling the genome as a circular construct may be more efficient. Since the genome forms concatemers during the normal replication cycle, we also reasoned that the circularized genome could be rebooted in *E coli* (Kulczyk & Richardson 2016). This approach is also how the genome would be assembled into a plasmid backbone for further manipulation in another chassis (e.g., yeast), which seems worth exploring (Jaschke et al. 2012).

We designed primers with homology arms between the ends of the genome (fragments 1 and 6), creating an overlapping junction and making a circular product upon assembly. After confirmation of the successful assembly of each junction **(Figure 5B)**, we transformed the Gibson Assembly mixture to see if any plaques could form. Unexpectedly, rebooting the circularly assembled genome seemed to yield fewer plaques than the linear assembly **(Figure 5A)**. It is not clear whether this is a result of reduced assembly efficiency *in vitro* or due to the less efficient rebooting of the circular genome. While the experiment yielded viable phage/plaques, it is not clear if the circular DNA yielded the plaques or if non-circularized DNA formed the plaques. Because the efficiency of using circular DNA assembly seemed much lower, we did not explore this further.

**Figure 5:**
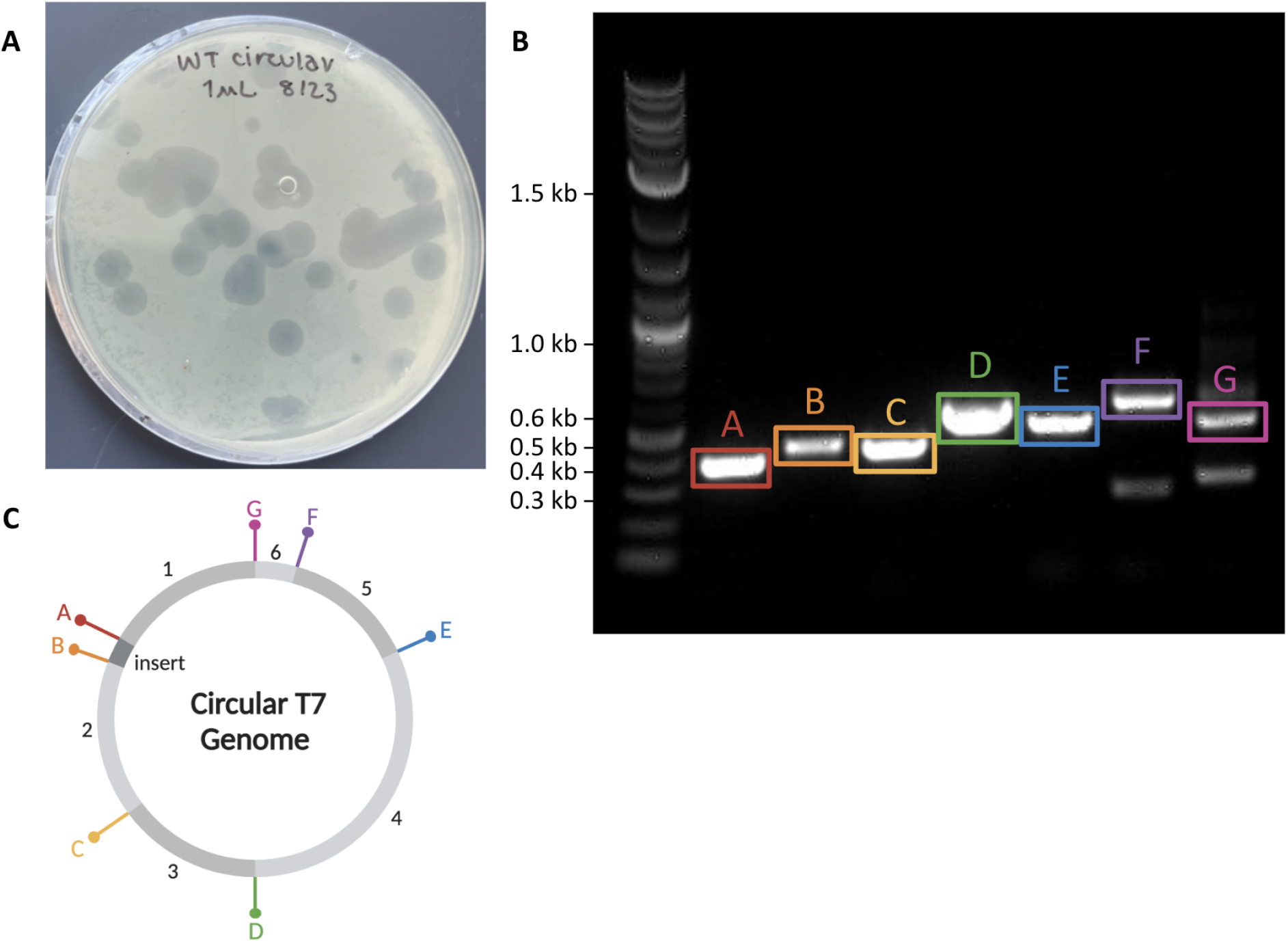
Circular assembly of phage containing landing pad insert. **(A)** The WT phage genome assembled as a circular DNA transformed into DH10B. **(B)** PCR assessment of *in vitro* circular assembly. Gibson Assembly were amplified via PCR using primers spanning the assembly junctions. **(C)** Genome map of circular assembled phage with location of respective junctions.

### Construction of the Genetic Landing Pad

To enable high-throughput screening of inserts, our goal was for SapI digestion to cut out the ‘landing pad’ (insertion region of interest) without modifying the rest of the genome. The landing pad region would then be prepared for DNA fragment insertion (a riboregulator, in this case) using GoldenGate assembly (Engler et al. 2008), Gibson Assembly (Gibson et al. 2009), or through ligation of compatible ends. To do so, we designed a landing pad at the region of interest flanked with inverted SapI sites, which would be removed along with the landing pad upon digestion. Along with the landing pad design, existing SapI sites were removed from the rest of the genome to prevent off-target digestion. These wild-type SapI sites were removed with mutations previously shown to produce bioactive phage (Pryor et al. 2022) **(Figure 1A)**. We then designed an “insert” to replace the landing pad at the ribosome binding site (RBS) upstream of gene 2.5. The RBS upstream of gene 2.5 was identified as a predicted hairpin in the WT T7 genome, which was confirmed by analyzing the sequences surrounding and including gene 2.5 with the *De Novo* DNA RBS Calculator (Predict Mode). **(Figure 6A)**.

**Figure 6:**
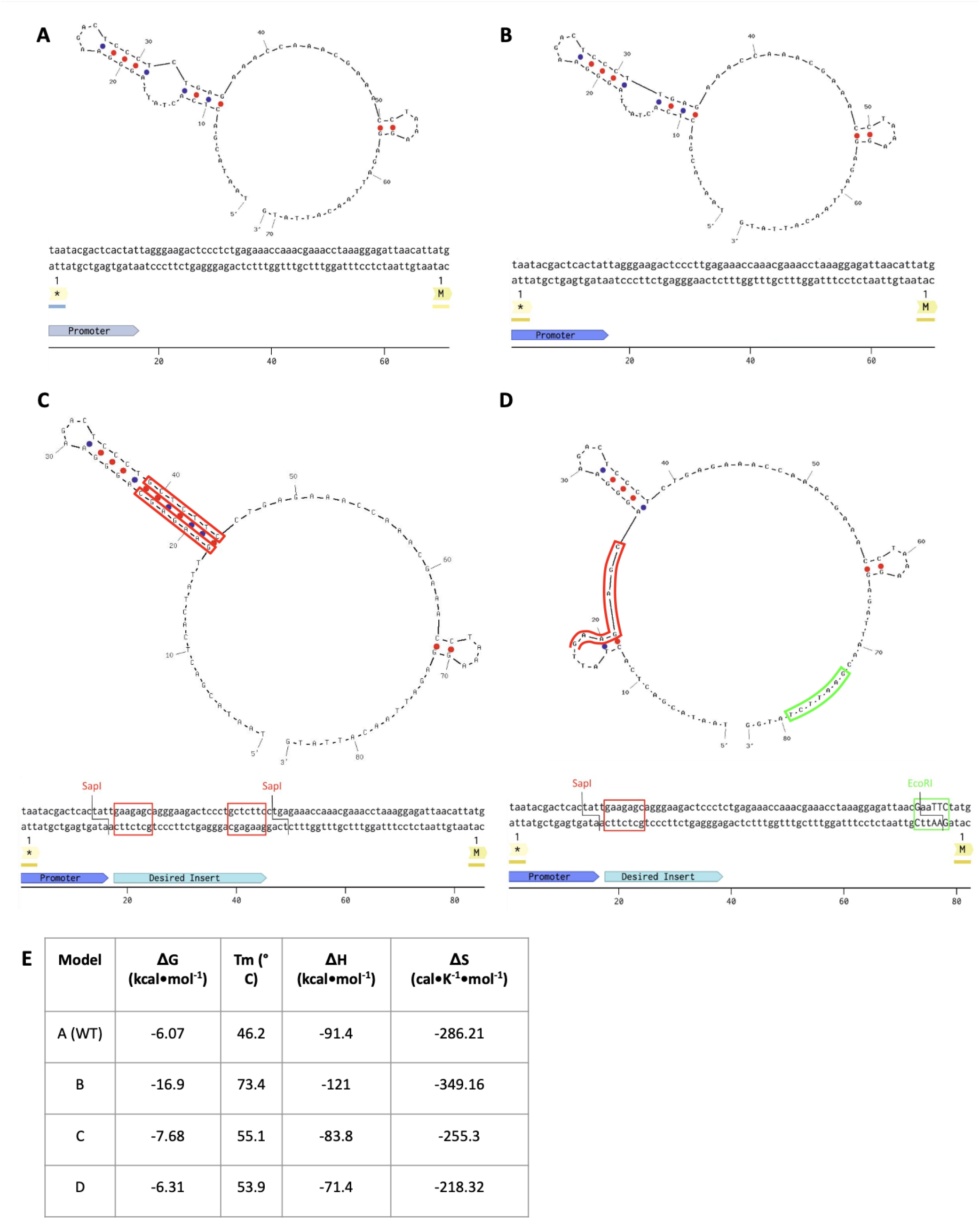
Dual Sapl containing landing pad alters mRNA structure and phage growth. The red and green box indicate the location of the recognition sequence of Sapl and respectively with respect to the landing pad insert. **(A)** Predicted hairpin formation of the WT sequence of gene 2.5 mRNA **(B)** Predicted hairpin formation of the mutated gene 2.5 mRNA following genome assembly and rebooting in *E. coli* does not contain Sapl sites. **(C)** Predicted hairpin formation of gene 2.5 mRNA containing inverted Sapl sites. **(D)** Predicted hairpin formation of gene 2.5 mRNA containing Sapl and **(E)** Table with predicted thermodynamic parameters for mRNA structures

Our initial attempts to reboot phage with two SapI sites consistently yielded very few and small phage plaques. Upon genomic sequencing of the phage recovered from these plaques, we found undesired mutations in the expected sequence of the landing pad, likely arising from recombination between the two SapI sites **(Figure 6B)**. We suspected that the two flanking SapI sites in opposite directions might enable the formation of a hairpin in front of the start codon of gene 2.5. Using IDT OligoAnalyzer, we found the flanking SapI sites, and the surrounding sequences formed a hairpin with a very high T_m_ (Zuker 2003) **(Figure 6C)**. Using the same analysis, we compared the sequence of the mutated phage and found both SapI sites had been mutated from the region **(Figure 6B, Figure 7)**, and the resulting hairpin looked similar to the WT sequence **(Figure 6A)**. We reasoned that the hairpin formed by the sequence of the two SapI sites inhibits translation, and that the only phage containing mutations where this hairpin was removed was able to grow. This validated our approach to regulating phage production via sequences upstream of gene 2.5. Because of this finding, we redesigned the landing pad to contain two different restriction sites.

**Figure 7:**
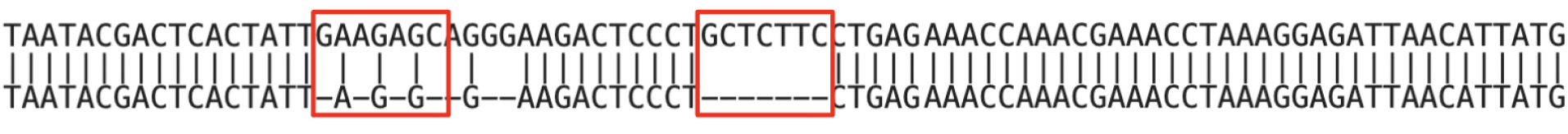
Alignment of WT phage (top) with phage that grew from transformed assemblies of the dual Sapl construct. The Sapl sites in the WT sequence are highlighted in red. The mutant that grew did not contain any Sapl sites.

The newly designed insert contained the native ribosome binding site flanked by non-native SapI and EcoRI restriction sites. The EcoRI restriction site was selected as it is one of the few commonly used restriction enzymes whose recognition sequence was not already found in the entire WT T7 genome. The SapI and EcoRI sites were positioned so that the introduced mutations required to create the recognition site utilized as many existing nucleotides from the native genomic sequence as possible, minimizing disruption to the reading frame. Designing a unique EcoRI site as part of the cloning site requires five base pairs of matching sequences. However, a similar strategy could be used with other non-Type IIS restriction sites to avoid recombination and make scarless cuts in a phage genome landing pad. Alternatively, two different Type IIS restriction sites could be used. We found a small hairpin in this new construct that matched the hairpin found in the WT sequence **(Figure 6A)**, which did not contain the SapI or EcoRI sites **(Figure 6D)**.

## Conclusion

Gibson Assembly is a simple and efficient way to assemble phage genomes from fragments. Unlike other approaches that treat fragments with an exonuclease and package the partially assembled fragments into phage particles *in vitro*, homolog-based assembly and direct transformation of the DNA do not require phage packaging or a TXTL kit. This makes assembly cheaper and more straightforward. Since Gibson Assembly does not rely on restriction enzymes, we were able to remove the SapI sites from the phage genome and use SapI inside our landing pad. Gibson Assembly and other homology-based assemblies allow for this flexibility, which is necessary when domestication interferes with the phage function of the genome design.

In our experiments, this flexibility allowed us to easily optimize the landing pad upstream of gene 2.5. In doing this, we were able to determine that the sequences upstream of gene 2.5 of the T7 phage were critical, making a good target for an RNA switch to modulate phage replication. Similar approaches could be taken to control other phages by installing specific sequences upstream of an essential gene. By using a landing pad, only a single fragment would need to be inserted into the digested genome, the final assembly would comprise only three fragments. This should be much more robust than the assembly of the genome from multiple fragments, as we initially did to produce the genome. Reassembly after digesting the landing pad can be achieved by Gibson Assembly or other methods, including Golden Gate, where the fragment to be inserted is flanked by Type IIS restriction sites (assuming they do not interfere with function). Based on our results, using restriction sites with two different sequences can minimize issues with the hairpin formation of the mRNA. If Golden Gate was used to assemble the genome with the landing pad, the restriction enzyme used for Golden Gate could not be engineered into the landing pad itself. Hence, if two different Type IIS sites were used in the landing pad, the genome would have to have both restriction sites removed in addition to a third one for assembling with Golden Gate. Once Gibson Assembly is used to construct the genome with the landing pad, Golden Gate may be used to insert fragments into the landing site by simply combining the phage DNA with the insert or with a library of DNAs.

It still remains to be seen if Gibson Assembly or packaging the exonuclease-treated DNA is more efficient. In studies using either method, the number of fragments and overlap length varied, and the optimal overlap size may differ between the two methods. Since Gibson Assembly works well with many fragments and shorter homology arms than was done with a packaging approach (Nozaki 2022), and since optimizing Gibson Assembly is fairly standardized, this approach is more accessible, even if it is less efficient. In addition, Gibson Assembly reactions can be troubleshot or optimized using diagnostic PCRs to assess which junctions form and which do not. Long read sequencing can also be used and provides additional insight into which junctions form, but to a lesser extent over junctions. This modular approach not only simplifies the process of genetic engineering in bacteriophages but also broadens the potential for their application in various biotechnological fields.

## Materials and Methods

### Strains

The Phage T7 wild-type (WT) genome was obtained from Boca Scientific Inc. (NC1143564). Synthetic phages were synthesized by PCR of the T7 WT genome to remove native SapI restriction sites and include an insert with flanking EcoRI and SapI site upstream of Gene 2.5. *Escherichia coli* strain DH10B was obtained from New England Biolabs (C3019I, obtained from New England Biolabs) and used to reboot modified phages. Phages were propagated in *E. coli* strain BL21 (C2530H, also obtained from New England Biolabs). For subcloning, an endonuclease I deficient *E. coli* strain, DH10B, was used to protect genome ends following transformation (Durfee et al. 2008). To verify the assembly, phages were propagated in *E. coli*, followed by DNA extraction and long-read sequencing.

### Culture Conditions

*E. coli* strains were grown in Luria Broth (LB) and 1.5% LB agar plates at 37°C. Phages were rebooted by electroporating an assembled phage genome into BL21 and grown in soft agar (0.6%) in LB on top of a solid LB agar medium at 37°C overnight before storage at 4°C.

### Generation of PCR Fragments for Phage Genome Assembly

Fragments of the WT T7 genome were amplified using PCR, utilizing KOD XtremeHot Start DNA Polymerase (Sigma-Aldrich, cat. #71975). Site-directed mutagenesis was utilized to remove native SapI sites from the WT T7 genome. Each adjacent DNA fragment had overlapping homology arms ≥ 30 bp **(Table 1)**. Due to mispriming events caused by 160 bp long direct repeats at both ends of the T7 genome, we gel purified each fragment was gel purified (in 1% agarose gel, cast with Invitrogen™ UltraPure™ Agarose-1000, (ThermoFisher Scientific, 16550100), utilizing the QIAquick Gel Extraction Kit (Qiagen 28704) and eluting in nuclease-free water from Zymo-Spin I Columns (Zymo Research C1003-250). Linear sequencing was performed by Plasmidsaurus for each T7 genome fragment using Oxford Nanopore Technology with custom analysis and annotation to confirm correct sequences. It is noteworthy that using KOD-Xtreme provides ample DNA yields following PCR and gel purification. This leads to highly pure DNA that can be easily sequenced and assembled.

**Table 1:**
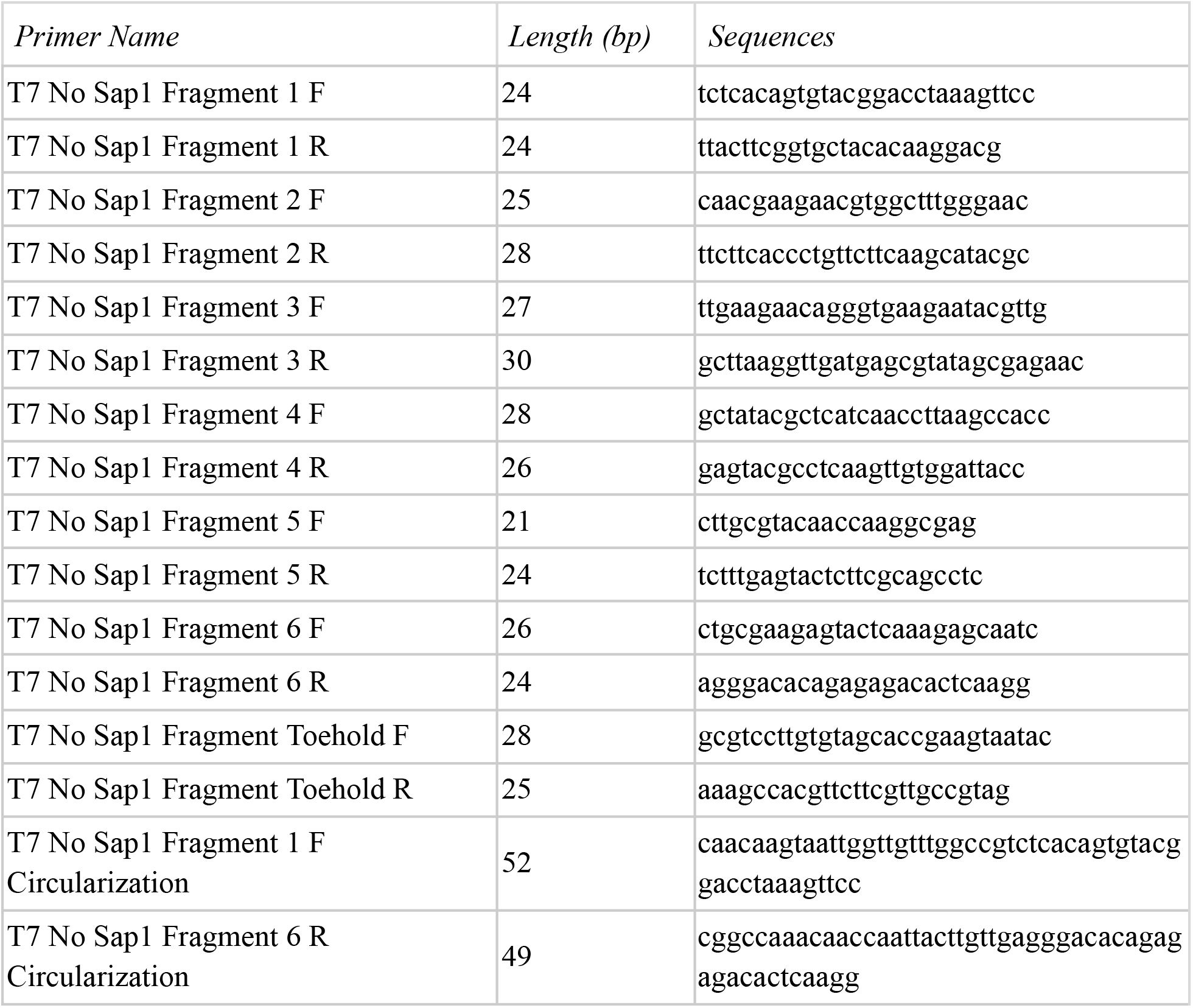
Primers used in this study for synthesis of fragments of the modified T7 phage genome.

### In Vitro Gibson Assembly of Phage Genome

The insert was synthesized with 30 bp homology to both adjacent T7 DNA fragments by Integrated DNA Technologies (Coralville, IA, USA). 0.05 pmol of each phage genome fragment, including the insert, was mixed with the NEBuilder HiFi DNA Assembly Master Mix (New England Biolabs, E2621X) and incubated at 50°C for 1one hour. The NEBuilder reaction mixture was heat-inactivated at 65°C for 20 minutes and PCR purified with a QIAquick PCR Purification Kit (Qiagen, 28104). The reaction product was submitted to Plasmidsaurus for long-read linear sequencing using Oxford Nanopore Technology.

### Verification of Assembled Phage Genome “Junctions”

We utilized PCR to verify successful assembly by designing primers **(Table 2)** to amplify a 300-500 bp region at the junction of each fragment from the Gibson Assembly mixture, employing the Invitrogen™ PCR Master Mix Starter Pack (A31021, ThermoFisher Scientific). The junction PCR reaction was visualized on 1% agarose gel, cast with Invitrogen™ UltraPure™ Agarose-1000 (16550100, ThermoFisher Scientific) against 1 kb+ DNA ladder (N3200S, New England Biolabs), to quickly confirm successful assembly prior to a plaque assay.

**Table 2:**
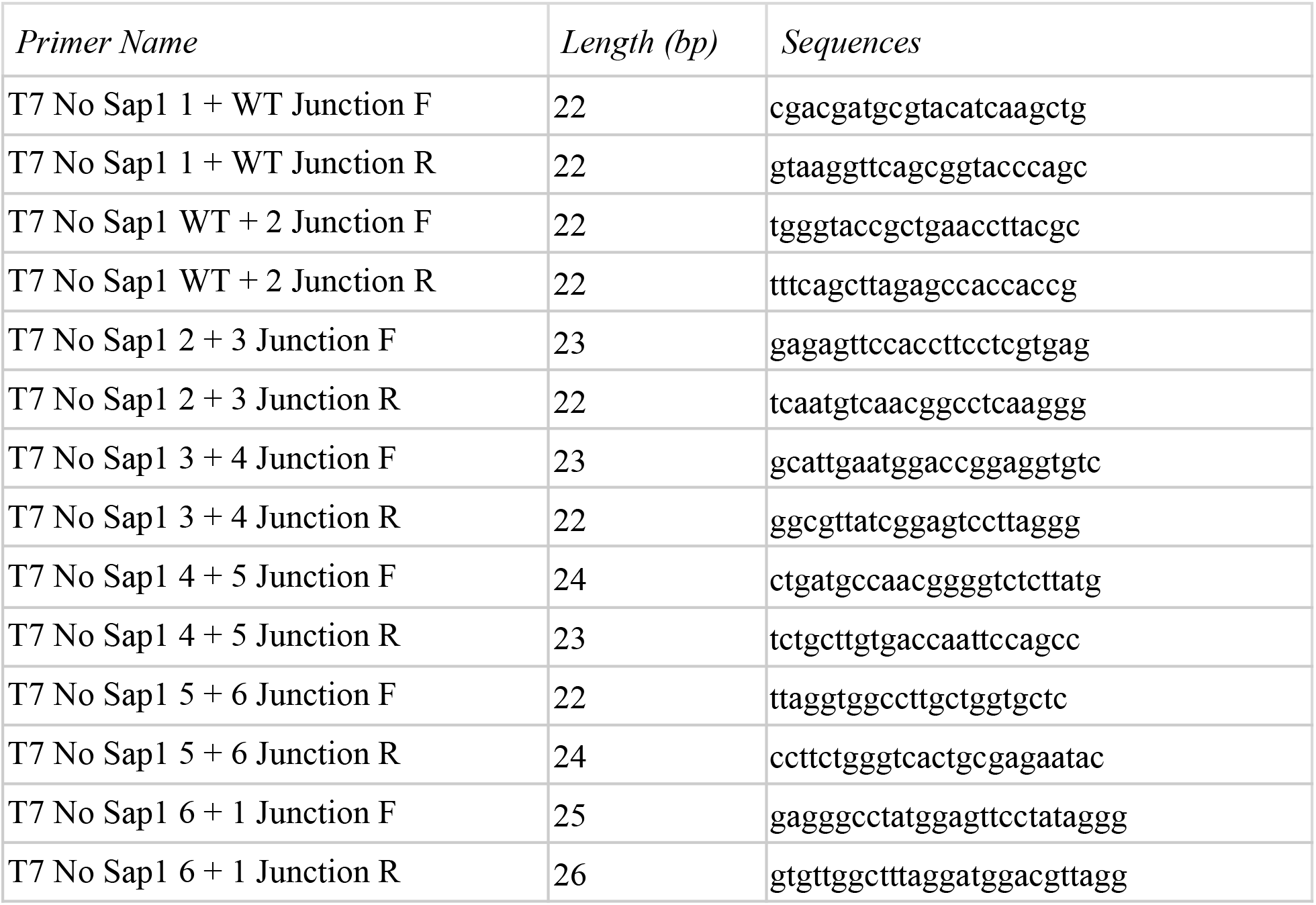
Primers used in this study for the amplification of assembled fragment junctions.

### Modified Phage Rebooting

To reboot assembled phages, 1 µL of assembly mixture was electroporated into 50 µL of DH10B cells in a 1 mm electrode spacing cuvette (Bio-Rad) and a Gene Pulser Xcell Electroporation System (Bio-Rad) at the following settings: 1.40 kV voltage, 50 F capacitance, and 150 Ω resistance. Cells were recovered at 37°C for 15 min in 500 µL of SOC medium and mixed into 3 mL 0.6% soft agar, pre-warmed to 55°C. The mixture was poured into pre-warmed LB plates and incubated overnight at 37°C. Plaque formation indicated successfully assembled phage genomes, as partially assembled genomes would not yield plaques.

### Rebooted Phage Propagation

To propagate the rebooted phage, 100 mL LB was spiked with 0.1 volumes of overnight *Escherichia coli* strain BL21 culture and incubated with agitation for one hour at 37°C. A biopsy punch was taken from a plaque of the rebooted phage and added to the bacterial host culture. The culture was incubated with agitation at 37°C and 275 rpm for ~5 hours until the lysate cleared.

Phage lysate was aliquoted into sterile 50 mL centrifuge Falcon tubes and centrifuged at 4,000 g for 20 minutes at RT. Supernatant was collected and filter-sterilized using a 0.22 µm filter to yield a bacterial cell-free phage lysate. The 5×phage precipitation solution (20% w/v PEG 8000, 2.5 M NaCl) was added to the lysate and incubated overnight at 4°C. Lysate was centrifuged at 10,000×g 4°C for 30 min. The precipitated phage pellet was resuspended in 1× phage resuspension buffer (1M NaCl, 10 mM Tris•HCl pH 7.5, 0.1 mM EDTA). 10× DNAse I buffer (10 mM Tris•HCl pH 7.6, 2.5 mM MgCl_2_, 0.5 mM CaCl_2_) was added to final 1× concentration and mixed by gently vortexing. 1 µL DNAse I (2,000 U/mL), obtained from New England Biolabs (M0303), and 1 µL RNase A (20 mg/mL), obtained from Invitrogen PureLink™ #12091021, was added to the resuspended phage pellet and incubated for 30 minutes at 37°C to degrade remaining nucleic acids and lysed bacterial cells. 10 µL of 0.5 M EDTA (pH 8.0) was added to chelate divalent metals.

### Rebooted Phage Phenol:Chloroform DNA Extraction

The rebooted phage DNA was extracted from purified phage particles using phenol:chloroform:isoamyl alcohol (25:24:1) method and precipitated using ethanol (100%). One volume of phenol:chloroform:isoamyl alcohol (25:24:1) was added to resuspended phage precipitation, and the content was centrifuged at 16,000×g for 5 minutes. The aqueous phase was extracted into a new tube, and 0.1 volume of 3 M sodium acetate and 2.5 volumes 100% ethanol, prechilled to –20°C, was added. The sample was incubated at –20°C for 30 minutes to overnight. The solution was centrifuged at 14,000×g 4°C for 15 minutes, and the supernatant was aspirated. The pellet was washed with 500 µL of 70% ethanol, pre-chilled at –20°C, and air dried. The DNA pellet was resuspended in 10 mM Tris•HCl pH 7.5 and stored at –20°C.

The purified phage DNA was visualized on 1% agarose gel, and the long-read whole genome sequencing was performed by Plasmidsaurus using Oxford Nanopore Technology with custom analysis and annotation. This experiment was repeated eight times with rebooted phage plaques to verify successfully edited phage genomes.

## Acknowledgements

We would like to express our sincere gratitude for the support and resources that made this research possible. We acknowledge making@Stanford and the Biological Interdisciplinary Open Maker Environment (BIOME) for their funding and the Uytengsu Teaching Lab (UTL) for providing the essential lab space for our experiments. We are thankful to the Stanford University Bioengineering Department for their continuous support throughout this project. This research was conducted under the auspices of iGEM, and we would like to acknowledge the contributions of the Stanford iGEM Team, whose collaborative efforts and dedication were instrumental in the completion of this work. Plasmidsaurus for free long-read sequencing.

## Supporting Files

https://github.com/juliavu2004/phage

