## Supplementary material for "Homology-Based Enzymatic Assembly of Modular T7 Phage Genome": Table 1_ Primers used in this study for synthesis of fragments of the modified T7 phage genome.docx

| Primer Name | Length (bp) | Sequences |
| --- | --- | --- |
| T7 No Sap1 Fragment 1 F | 24 | tctcacagtgtacggacctaaagttcc |
| T7 No Sap1 Fragment 1 R | 24 | ttacttcggtgctacacaaggacg |
| T7 No Sap1 Fragment 2 F | 25 | caacgaagaacgtggctttgggaac |
| T7 No Sap1 Fragment 2 R | 28 | ttcttcaccctgttcttcaagcatacgc |
| T7 No Sap1 Fragment 3 F | 27 | ttgaagaacagggtgaagaatacgttg |
| T7 No Sap1 Fragment 3 R | 30 | gcttaaggttgatgagcgtatagcgagaac |
| T7 No Sap1 Fragment 4 F | 28 | gctatacgctcatcaaccttaagccacc |
| T7 No Sap1 Fragment 4 R | 26 | gagtacgcctcaagttgtggattacc |
| T7 No Sap1 Fragment 5 F | 21 | cttgcgtacaaccaaggcgag |
| T7 No Sap1 Fragment 5 R | 24 | tctttgagtactcttcgcagcctc |
| T7 No Sap1 Fragment 6 F | 26 | ctgcgaagagtactcaaagagcaatc |
| T7 No Sap1 Fragment 6 R | 24 | agggacacagagagacactcaagg |
| T7 No Sap1 Fragment Toehold F | 28 | gcgtccttgtgtagcaccgaagtaatac |
| T7 No Sap1 Fragment Toehold R | 25 | aaagccacgttcttcgttgccgtag |
| T7 No Sap1 Fragment 1 F Circularization | 52 | caacaagtaattggttgtttggccgtctcacagtgtacggacctaaagttcc |
| T7 No Sap1 Fragment 6 R Circularization | 49 | cggccaaacaaccaattacttgttgagggacacagagagacactcaagg |

Table 2: Primers used in this study for amplification of assembled fragment junctions.

| Primer Name | Length (bp) | Sequences |
| --- | --- | --- |
| T7 No Sap1 1 + WT Junction F | 22 | cgacgatgcgtacatcaagctg |
| T7 No Sap1 1 + WT Junction R | 22 | gtaaggttcagcggtacccagc |
| T7 No Sap1 WT + 2 Junction F | 22 | tgggtaccgctgaaccttacgc |
| T7 No Sap1 WT + 2 Junction R | 22 | tttcagcttagagccaccaccg |
| T7 No Sap1 2 + 3 Junction F | 23 | gagagttccaccttcctcgtgag |
| T7 No Sap1 2 + 3 Junction R | 22 | tcaatgtcaacggcctcaaggg |
| T7 No Sap1 3 + 4 Junction F | 23 | gcattgaatggaccggaggtgtc |
| T7 No Sap1 3 + 4 Junction R | 22 | ggcgttatcggagtccttaggg |
| T7 No Sap1 4 + 5 Junction F | 24 | ctgatgccaacggggtctcttatg |
| T7 No Sap1 4 + 5 Junction R | 23 | tctgcttgtgaccaattccagcc |
| T7 No Sap1 5 + 6 Junction F | 22 | ttaggtggccttgctggtgctc |
| T7 No Sap1 5 + 6 Junction R | 24 | ccttctgggtcactgcgagaatac |
| T7 No Sap1 6 + 1 Junction F | 25 | gagggcctatggagttcctataggg |
| T7 No Sap1 6 + 1 Junction R | 26 | gtgttggctttaggatggacgttagg |
